# Virome composition in marine fish revealed by meta-transcriptomics

**DOI:** 10.1101/2020.05.06.081505

**Authors:** Jemma L. Geoghegan, Francesca Di Giallonardo, Michelle Wille, Ayda Susana Ortiz-Baez, Vincenzo A. Costa, Timothy Ghaly, Jonathon C. O. Mifsud, Olivia M. H. Turnbull, David R. Bellwood, Jane E. Williamson, Edward C. Holmes

## Abstract

Revealing the determinants of virome composition is central to placing disease emergence in a broader evolutionary context. Fish are the most species-rich group of vertebrates and so provide an ideal model system to study the factors that shape virome compositions and their evolution. We characterised the viromes of 19 wild-caught species of marine fish using total RNA sequencing (meta-transcriptomics) combined with analyses of sequence and protein structural homology to identify divergent viruses that often evade characterisation. From this, we identified 25 new vertebrate-associated viruses and a further 22 viruses likely associated with fish diet or their microbiomes. The vertebrate-associated viruses identified here included the first fish virus in the *Matonaviridae* (single-strand, negative-sense RNA virus). Other viruses fell within the *Astroviridae, Picornaviridae, Arenaviridae, Reoviridae, Hepadnaviridae, Paramyxoviridae, Rhabdoviridae, Hantaviridae, Filoviridae* and *Flaviviridae* and were sometimes phylogenetically distinct from known fish viruses. We also show how key metrics of virome composition – viral richness, abundance and diversity – can be analysed along with host ecological and biological factors as a means to understand virus ecology. Accordingly, these data suggest that that the vertebrate-associated viromes of the fish sampled here are predominantly shaped by the phylogenetic history (i.e. taxonomic order) of their hosts, along with several biological factors including water temperature, habitat depth, community diversity and swimming behaviour. No such correlations were found for viruses associated with porifera, molluscs, arthropods, fungi and algae, that are unlikely to replicate in fish hosts. Overall, these data indicate that fish harbour particularly large and complex viromes and the vast majority of fish viromes are undescribed.

## Introduction

Metagenomic next-generation sequencing (mNGS) has led to a revolution in virus discovery (Shi et al 2018b, Zhang et al 2018a, Zhang et al 2018b), exposing more of the diversity, scale and structure of the virosphere. However, while it is now possible to reveal host viromes en masse (Chang et al 2019, Vibin et al 2018, Geoghegan et al 2018b, Paez-Espino et al 2016, Lim et al (2015), Porter et al 2019, Roux et al 2017, Shi et al 2016, Temmam et al 2016, Tirosh et al 2018, Pettersson et al 2019), we still have an incomplete understanding of the factors that structure viromes. Until recently, studies of virus evolution were largely limited to single viruses and/or single hosts, restricting our ability to explore the diverse host and environmental factors that might structure viromes as a whole. Fortunately, this is changing with the advent of mNGS, particularly total RNA sequencing. In particular, metagenomic-based studies have shown that aspects of host biology can greatly impact virus diversification (Wille et al 2019, Wille 2020) and as such may also be key drivers of virus emergence. As a simple case in point, the behavioural ecology of host species directly affects contact rates among individuals in a population, and more frequent intra- and inter-species contacts are likely to increase the potential for viral transmission.

The marine environment is a rich source of viruses. For example, the bacteriophage in aquatic ecosystems greatly outnumber other life-forms (Maranger and Bird 1995). There is an estimated concentration of 10 billion virus particles per litre of surface water (Bergh et al 1989, Breitbart and Rohwer 2005, Middelboe and Brussaard 2017, Suttle 2005), although abundance levels vary with such factors as ocean depth (De Corte et al 2012, Lara et al 2017), temperature (Coutinho et al 2017), latitude (Gregory et al 2019) and phytoplankton bloom development (Alarcon-Schumacher et al 2019). In marked contrast to bacteriophage, little is known about the factors that contribute to virus diversity in aquatic vertebrate populations, even though viruses can cause large-scale disease outbreaks in farmed fish (Crane and Hyatt 2011, Jarungsriapisit et al 2020, Whittington and Reddacliff 1995).

Fish provide an ideal model to better understand the diversity of viruses that exist in nature as well as the range of host and environmental factors that shape virome composition and abundance. Fish are the most species-rich group of vertebrates with over 33,000 species described to date (fishbase.org), the vast majority of which (∼85%) are bony fish (the Osteichthyes) (Betancur-R et al 2017). Bony fish themselves are an extremely diverse and abundant group comprising 45 taxonomic orders, exhibiting a wide range of biological features that likely play an important role in shaping the diversity of their viromes. Initial studies indicate that fish harbour a remarkable diversity of viruses, particularly those with RNA genomes, that may exceed that seen in any other class of vertebrate (Geoghegan et al 2018a, Lauber et al 2017, Shi et al 2018a). In addition, those viruses present in fish often appear to be the evolutionary predecessors of viruses infecting other vertebrate hosts, generally indicative of a pattern of virus-host associations that can date back hundreds of millions of years, although with frequent cross-species transmission. Despite the apparent diversity and ubiquity of fish viruses, they are severely under-studied compared to mammalian and avian viruses and there is little data on the factors that determine the structure of fish viromes.

To reveal more of the unexplored aquatic virosphere we sampled wild-caught ray-finned marine fish spanning 23 species across nine taxonomic orders and quantified a variety of host characteristics that together may impact virome composition, abundance and evolution. Specifically, we utilised meta-transcriptomics together with both sequence and protein structural homology searches of known viruses to: (i) reveal the total virome composition of fish, (ii) describe the phylogenetic relationships of the novel viruses obtained, (iii) determine whether, on these data, there may be associations between virome composition, abundance, richness and diversity and particular host traits, and (iv) explore whether taxonomically-related fish hosts have more similar viromes. The host characteristics initially considered here were: fish taxonomic order, swimming behaviour (i.e. solitary or schooling fish), preferred climate, mean preferred water temperature, host community diversity (i.e. multi- or single-species community), average body length, maximum life span, trophic level, and habitat depth (SI Table 1).

## Methods

### Ethics

Biosafety was approved by Macquarie University, Australia (ref: 5201700856). This study involved dead fish purchased from a fish market for which no animal ethics approval was required. The pygmy goby was collected under GBRMPA permit G16/37684.1 and JCU Animal Ethics Committee #A2530.

### Fish sample collection

Dead fish from 23 species were sampled for virome analysis (SI Table 1). These included 18 new species collected from a fish market in Sydney, Australia, together with four species from our previous sampling of the same fish market (Geoghegan et al 2018a). These animals were caught by commercial fisheries in coastal waters in New South Wales, Australia by several different suppliers in Autumn 2018. By way of contrast, an additional species, the pygmy goby (*Eviota zebrina*), was obtained from the coral reefs of tropical northern Queensland at approximately the same time. Fish were snap frozen at - 20°C immediately upon capture. Fish obtained from the market were purchased on the day of catch. Tissues were dissected and stored in RNALater before being transferred to a - 80°C freezer. To increase the likelihood of virus discovery during metagenomic sequencing, 10 individuals from each species were pooled.

### Transcriptome sequencing

mNGS was performed on fish tissue (liver and gill). Frozen tissue was partially thawed and submerged in lysis buffer containing 1% ß-mercaptoethanol and 0.5% Reagent DX before tissues were homogenized together with TissueRupture (Qiagen). The homogenate was centrifuged to remove any potential tissue residues, and RNA from the clear supernatant was extracted using the Qiagen RNeasy Plus Mini Kit. RNA was quantified using NanoDrop (ThermoFisher) and tissues from each species were pooled to 3μg per pool (250ng per individual). Libraries were constructed using the TruSeq Total RNA Library Preparation Protocol (Illumina) and host ribosomal RNA (rRNA) was depleted using the Ribo-Zero-Gold Kit (Illumina) to facilitate virus discovery. Paired-end (100bp) sequencing of the RNA library was performed on the HiSeq 2500 platform (Illumina). All library preparation and sequencing were carried out by the Australian Genome Research Facility (AGRF).

### Transcript sequence similarity searching for viral discovery

Sequencing reads were first quality trimmed then assembled *de novo* using Trinity RNA-Seq (Haas et al 2013). The assembled contigs were annotated based on similarity searches against the NCBI nucleotide (nt) and non-redundant protein (nr) databases using BLASTn and Diamond (BLASTX) (Buchfink et al 2015), and an e-value threshold of 1×10^−5^ was used as a cut-off to identify positive matches. We removed non-viral hits including host contigs with similarity to viral sequences (e.g. endogenous viral elements). To reduce the risk of incorrect assignment of viruses to a given library due to index-hoping, those viruses with a read count less than 0.1% of the highest count for that virus among the other libraries was assumed to be contamination.

### Protein structure similarity searching for viral discovery

To identify highly divergent viral transcripts, particularly those that might be refractory to detection using similarity searching methods such as the BLAST approach described above, we employed a protein structure-based similarity search for ‘orphan’ contigs that did not share sequence similarity with known sequences. Accordingly, assembled orphan contigs were translated into open reading frames (ORFs) using EMBOSS getorf program (Rice et al 2000). ORFs were arbitrarily defined as regions between two stop codons with a minimum size of 200 amino acids in length. To reduce redundancy, amino acid sequences were grouped based on sequence identity using the CD-HIT package v4.6.5 (Li and Godzik 2006). The resulting data set was then submitted to Phyre2, which uses advanced remote homology detection methods to build 3D protein models, predict ligand binding sites, and analyse the effect of amino acid variants (Kelley et al 2015). Virus sequences with predicted structures were selected on the basis of having confidence values ≥90%. Following structure prediction, we used the associated annotations for preliminary taxonomic classification. To avoid false positives due to the limited number of available structures in the Protein Data Bank (PDB) for template modelling, the taxonomic assignment was cross-validated with the results from the Diamond (BLASTX) similarity search. Subsequently, putative viruses were aligned with reference viral protein sequences at the immediate higher taxonomic level (e.g. genus, family), using MAFFT v7.4 (E-INS-i algorithm) (Katoh and Standley 2013). Finally, we verified the similarity among sequences by careful visual inspection of the most highly conserved motifs of target proteins.

### Inferring the evolutionary history of fish viruses

We inferred the evolutionary relationships of the viruses contained in the fish samples and compared them with known viruses to determine those that were likely associated with vertebrate or non-vertebrate hosts. Specifically, we assumed that viruses that grouped with other vertebrate viruses in phylogenetic trees were likely to infect the fish sampled here, while those virus that were more closely related to those usually associated with other host types (such as invertebrates, fungi and plants) were unlikely to infect and replicate in fish hosts. To achieve this, the translated viral contigs were combined with representative protein sequences within each virus family obtained from NCBI RefSeq. The sequences retrieved were then aligned with those generated here again using MAFFT v7.4 (E-INS-i algorithm) as described above. Ambiguously aligned regions were removed using trimAl v.1.2 (Capella-Gutierrez et al 2009). To estimate phylogenetic trees, we selected the optimal model of amino acid substitution identified using the Bayesian Information Criterion as implemented in Modelgenerator v0.85 (Keane et al 2006) and analysed the data using the maximum likelihood approach available in IQ-TREE (Nguyen et al 2015) with 1000 bootstrap replicates. Phylogenetic trees were annotated with FigTree v.1.4.2. Viruses newly identified here were named reflecting the host common name.

### Revealing virome abundance and diversity

Transcriptomes were quantified using RNA-Seq by Expectation-Maximization (RSEM) as implemented within Trinity (Li and Dewey 2011). We first estimated the relative abundance of a host reference gene, ribosomal protein S13 (RPS13), to assess the sequencing depth across libraries. Next, we used RSEM to estimate the relative abundance of each virus transcript in these data.

For those viruses most likely associated with fish themselves, rather than components of their diet or microbiome (see Results), we performed analyses of virome abundance and diversity using R v3.4.0 integrated into RStudio v1.0.143 and plotted using ggplot2. Both the observed virome richness and Shannon effective (i.e. alpha diversity) were calculated for each library at the virus family level using modified Rhea script sets (Lagkouvardos et al 2017, Wille et al 2019). We used generalized linear models (GLM) to initially evaluate the effect of host taxonomic order, swimming behaviour (solitary or schooling fish), preferred climate, mean preferred water temperature, host community diversity, average species length, trophic level and habitat depth on viral abundance and alpha diversity (see SI Table 1 for all variables). Models were χ^2^ tested (LRT) to assess model significance. When the number of factor levels in an explanatory variable exceeded two, we conducted Tukey posthoc testing (glht) using the *multcomp* package (Hothorn et al 2008). Beta diversity (i.e. the diversity between samples) was calculated using the Bray Curtis dissimilarity matrix. Effects of variables on viral community composition were evaluated using permanova (Adonis Tests) and Mantel tests with 10,000 permutations using the *vegan* package (Oksanen 2007).

To establish connectivity (i.e. sharing) among virus families that were likely associated with non-fish hosts, we generated a cord diagram by quantifying the number of fish species harbouring each virus family identified in this study. Virus families that occur in the same fish species were represented by ribbons or links in the diagram.

## Results

We used mNGS to characterise viral transcripts from 23 marine fish spanning nine taxonomic orders: 19 species from this current study together with four from our previous work (Geoghegan et al 2018a). We combined data from our previous fish sampling to expand our data set and to apply novel viral protein structural searching methods not used previously. For these reasons, individual viruses discovered in our previous study are not detailed here. Combined, the extracted total RNA was organised into 23 libraries for high-throughput RNA sequencing. Ribosomal RNA-depleted libraries resulted in a median of 45,690,996 (range 33,344,520 – 51,071,142) reads per pool.

### Diversity and abundance of viruses in fish

The fish viromes characterised here contained viruses that were associated with vertebrate hosts as well as those that were more likely associated with porifera, invertebrates, fungi and algae (Figure 1). We primarily focused on the former since we assumed that the vertebrate-associated viruses were directly infecting the fish sampled, rather than being associated with the aquatic environment, diet or a co-infecting parasite, and hence are more informative in determining how host factors shape virus ecology and evolution.

**Figure 1.**
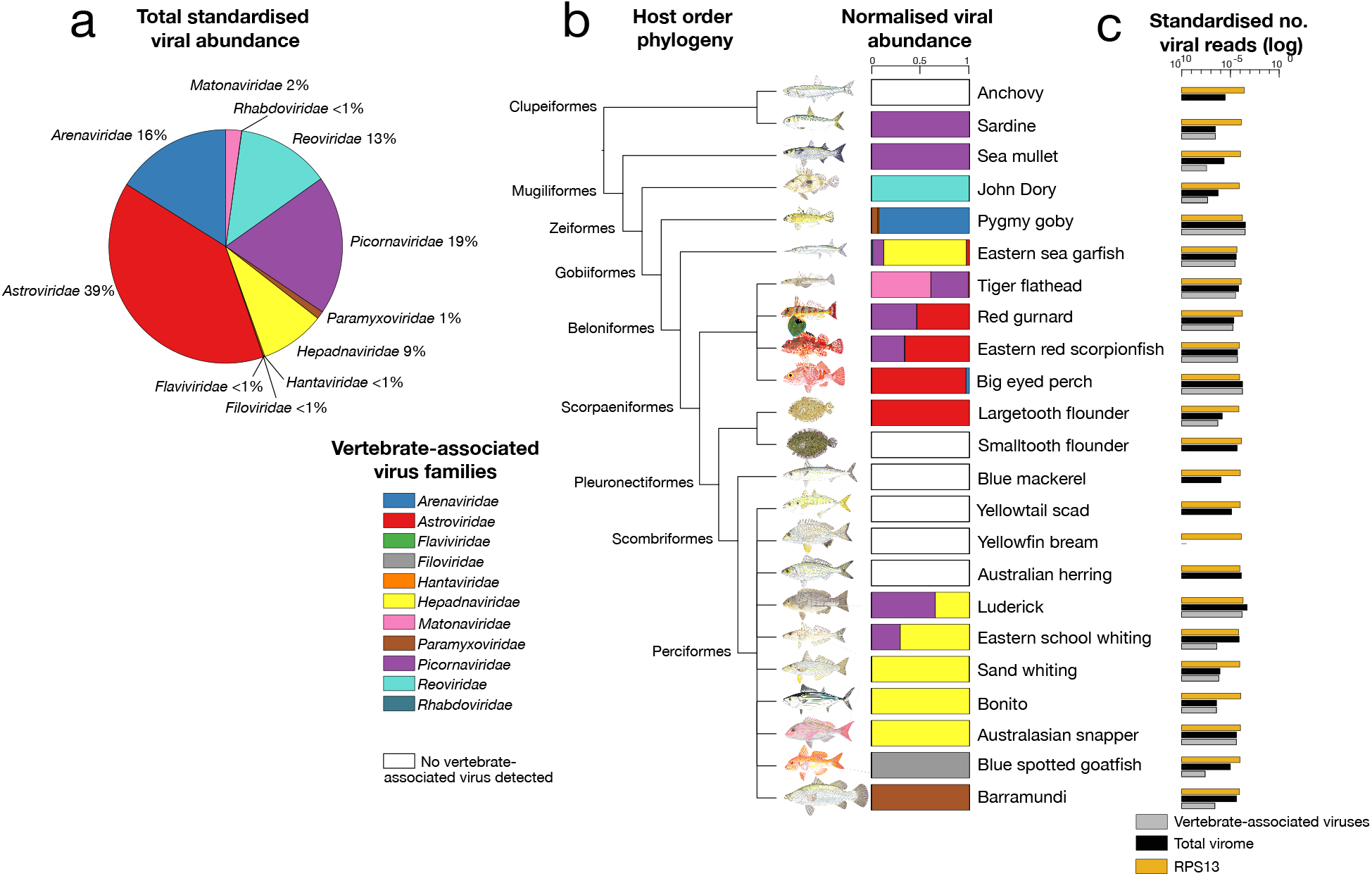
(A) Total standardized abundance of vertebrate-associated viruses (at the level of virus family) across the fish species examined. (B) Normalised viral abundance set out on a backbone of the fish host phylogeny at the order level. (C) Standardised number of total viral reads (black), vertebrate-associated viral reads (grey) and host reference gene ribosomal protein S13 (RPS13) (orange) in each species library.

Overall, we identified virus transcripts likely associated with vertebrate hosts that could be assigned to 11 viral families and present in a variety of fish species (SI Figure 1a). With the exception of the *Hepadnaviridae*, all were RNA viruses. Across all the fish sampled, those viral families found at relatively high abundances included the *Astroviridae* (representing 39% of all viruses discovered), *Picornaviridae* (19%), *Arenaviridae* (16%), *Reoviridae* (13%) and the *Hepadnaviridae* (9%) (Figure 1a). Other viral families found at lower relative abundances were the *Matonaviridae* (previously the *Togaviridae*) (2%), *Paramyxoviridae* (1%), as well as the *Rhabdoviridae, Hantaviridae, Filoviridae* and *Flaviviridae* (all <1%) (Figure 1a). The most common vertebrate-associated viruses found in these fish were picornaviruses (eight species), astroviruses (seven species) and hepadnaviruses (six species) (Figure 1b). The eastern sea garfish (*Hyporhamphus australis*) harboured the most diverse virome with four distinct vertebrate-associated viruses (Figure 1b). Six fish contained no vertebrate-associated viruses, and we found no viral sequences in the yellowfin bream (*Acanthopagrus australis*) (Figure 1c). An equivalent analysis of a host reference gene, ribosomal protein S13 (RPS13) that is stably expressed in fish, revealed similar abundances across species (0.004% – 0.02%), implying similar sequencing depth across libraries (Figure 1c). RPS13 was, on average, ∼55% more abundant than the total virome.

We also examined viruses that were phylogenetically related to those associated with porifera, molluscs, arthropods, fungi and algae, and hence were unlikely to infect the fish themselves. Accordingly, we identified an additional 22 viruses across 11 virus families (SI Figure 1b). These viruses were found in the *Chuviridae, Hepeviridae, Narnaviridae, Nodaviridae, Partitiviridae, Picornaviridae, Solemoviridae, Tombusviridae, Totiviridae, Dicistroviridae* and *Iflaviridae*, and are described in more detail below.

### Evolutionary relationships of fish viruses

To infer stable phylogenetic relationships among the viruses sampled and to identify those that are novel, where possible we utilised the most conserved (i.e. polymerase) viral regions that comprise the RNA-dependent RNA polymerase (RdRp) or the polymerase (P) ORF in the case of the hepadnaviruses. From this, we identified 25 distinct and potentially novel vertebrate-associated virus species, in addition to the eight novel viruses described previously (Geoghegan et al 2018a) (SI Table 2). All novel vertebrate-associated viruses shared sequence similarity to other known fish viruses with the exception of those viruses found in the *Matonaviridae* and *Rhabdoviridae*, the latter of which was found using structure similarity methods (Figure 2, SI Table 3; see below). We found a further 22 viruses that clustered with viruses found in porifera, molluscs, arthropods, fungi and algae (SI Figure 2, SI Figure 3, SI Figure 4).

**Figure 2.**
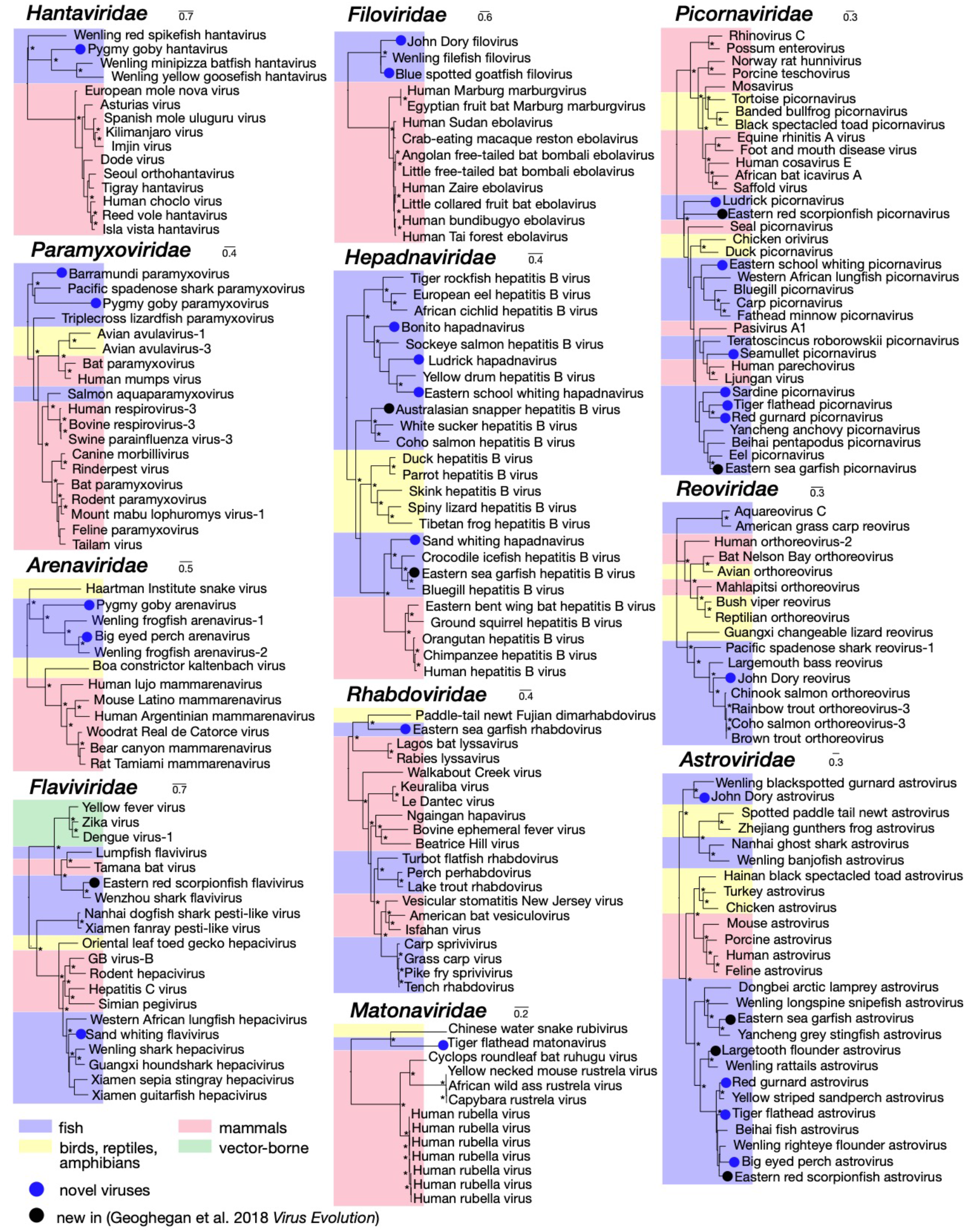
Phylogenetic relationships of likely vertebrate-associated viruses identified here. The maximum likelihood phylogenetic trees show the topological position of the newly discovered viruses (blue circles) and those identified in an earlier study (Geoghegan et al. 2018), in the context of their closest phylogenetic relatives. Branches are highlighted to represent host class (fish = blue; mammals = red; birds, reptiles and amphibians = yellow; vector-borne (mammals and arthropods) = green). All branches are scaled according to the number of amino acid substitutions per site and trees were mid-point rooted for clarity only. An asterisk indicates node support of >70% bootstrap support. See SI Table 3 for all accession numbers.

Among the viruses identified was tiger flathead matonavirus (in *Neoplatycephalus richardsoni*) – the first fish virus found in the *Matonaviridae*. This novel viral sequence shared only 35% amino acid similarity with its closest relative - Guangdong Chinese water snake rubivirus (Shi et al 2018a). Until recently, the only other representative of this family was the distantly related human rubella virus, although additional members of this family have recently been identified in other mammalian species (Bennett et al. 2020). Given the high levels of genetic divergence in this family, it is likely that these fish-associated viruses at least constitute a discrete and novel genus.

Another divergent virus discovered in this analysis is eastern sea garfish rhabdovirus (in *Hyporhamphus australis*), which was most closely related to Fujian dimarhabdovirus sampled from an amphibian host, sharing 45% amino acid RdRp sequence identity. Notably, this highly divergent virus was only identified by using protein structure homology, and forms a clade that is distinct from other fish rhabdoviruses (Figure 2). We also identified two novel viral sequences in the *Filoviridae* in John Dory (*Zeus faber*) and the blue spotted goatfish (*Upeneichthys lineatus*). These viruses shared sequence similarity to the only other known fish filovirus, Wenling filefish filovirus (Shi et al 2018a). With the exception of these fish viruses, all other known filoviruses including Ebola and Marburg viruses, are found in mammalian hosts, notably humans, bats and primates.

We also found numerous viruses that cluster within established clades of fish viruses. For example, pygmy goby hantavirus (in *Eviota zebrina*) grouped with other hantaviruses recently found in fish (Figure 2). Although they were previously only thought to infect mammals, hantaviruses have now been found to infect amphibians, jawless fish and ray-finned fish (Shi et al 2018a). The evolutionary history of the *Paramyxoviridae* shows two distinct fish virus lineages, of which both barramundi and pygmy goby paramyxoviruses grouped with Pacific spade-nose shark paramyxovirus and shared 50% and 45% amino acid L gene sequence similarity, respectively. This group of fish viruses is phylogenetically distinct from other paramyxoviruses. We also found novel fish viruses in the *Flaviviridae, Arenaviridae* and *Reoviridae*: although these grouped with other fish viruses, they greatly expand the known diversity of these virus families. Finally, as noted above, the most abundant viruses fell within the *Picornaviridae* and *Astroviridae*, and all shared sequence similarity to other fish viruses. Notably, both picornaviruses and astroviruses are single-stranded positive-sense RNA viruses that possess small icosahedral capsids with no external envelope, which may aid their preservation in harsh marine environments.

The only DNA viruses we identified were novel hepadnaviruses. Those found in bonito (*Sarda australis*), ludrick (*Girella tricuspidata*) and eastern school whiting (*Sillago flindersi*), fell into the divergent group of hepadna-like viruses, the nackednaviruses, that have been identified in a number of fish species (Lauber et al. 2017). In contrast, sand whiting hepadnavirus (in *Sillago ciliate*) fell into the fish virus clade that is more closely related to mammalian hepatitis B viruses (Dill et al 2016) (Figure 2).

As expected, many of the viruses identified here were associated with marine hosts belonging to invertebrates (including porifera, molluscs and arthropods; n = 20), fungi (n = 1) and algae (n = 1) as determined by their phylogenetic position and sequence similarity to viruses previously described in these taxa (SI Figure 2, SI Figure 3, SI Figure 4). This implies that these viruses more likely originated from host species that are associated with fish diet, fish microbiomes or the surrounding environment, rather than from the fish themselves. None of these viruses are highly divergent from other known viruses, but do help fill gaps in the phylogenetic diversity of these groups.

### Assessing the impact of host biology on virome composition

Our relatively small sample of 23 fish species precluded us from performing a detailed statistical analysis of the relationship between host traits and virome composition. Rather, we provide an initial analysis that should be regarded as a framework for understanding how key host variables might impact viral ecology and evolution, and that can be extended as more species are analysed.

To this end we examined the possible association between eight host traits and viral abundance (the proportion of viral reads in each sample), alpha diversity (the diversity within each sample, measured by observed richness and Shannon diversity) and beta diversity (the diversity between samples). The host traits initially considered here were: host taxonomic order, swimming behaviour (solitary or schooling fish), preferred climate, mean preferred water temperature, community diversity, average species length, maximum life span, trophic level and habitat depth.

We first focused on the vertebrate-associated virome. This initial analysis revealed that the phylogenetic relationships of the fish studied, as reflected in their taxonomic order, seemingly had the strongest association with the overall composition of fish viromes. This pattern was consistent when assessing viral abundance, alpha diversity and beta diversity (Figure 3). That is, fish order (χ^2^=0.003, df=8, p=0.0049) and mean preferred water temperature (χ^2^=0.008, df=1, p=0.035) were important predictors of viral abundance, such that Scopaeniformes (i.e. bigeye ocean perch, red gurnard, tiger flathead, and eastern red scorpionfish) had significantly higher viral abundance than Pleuronectiformes (i.e. largetooth and smalltooth flounder) (Tukey: z=3.766, p=0.00479), while viral abundance had a negative relationship to mean preferred water temperature (Figure 3). It is worth noting, however, that virus abundance within the Scopaeniformes were widely distributed and that their overall high abundance might only be due to a few species or individuals.

**Figure 3.**
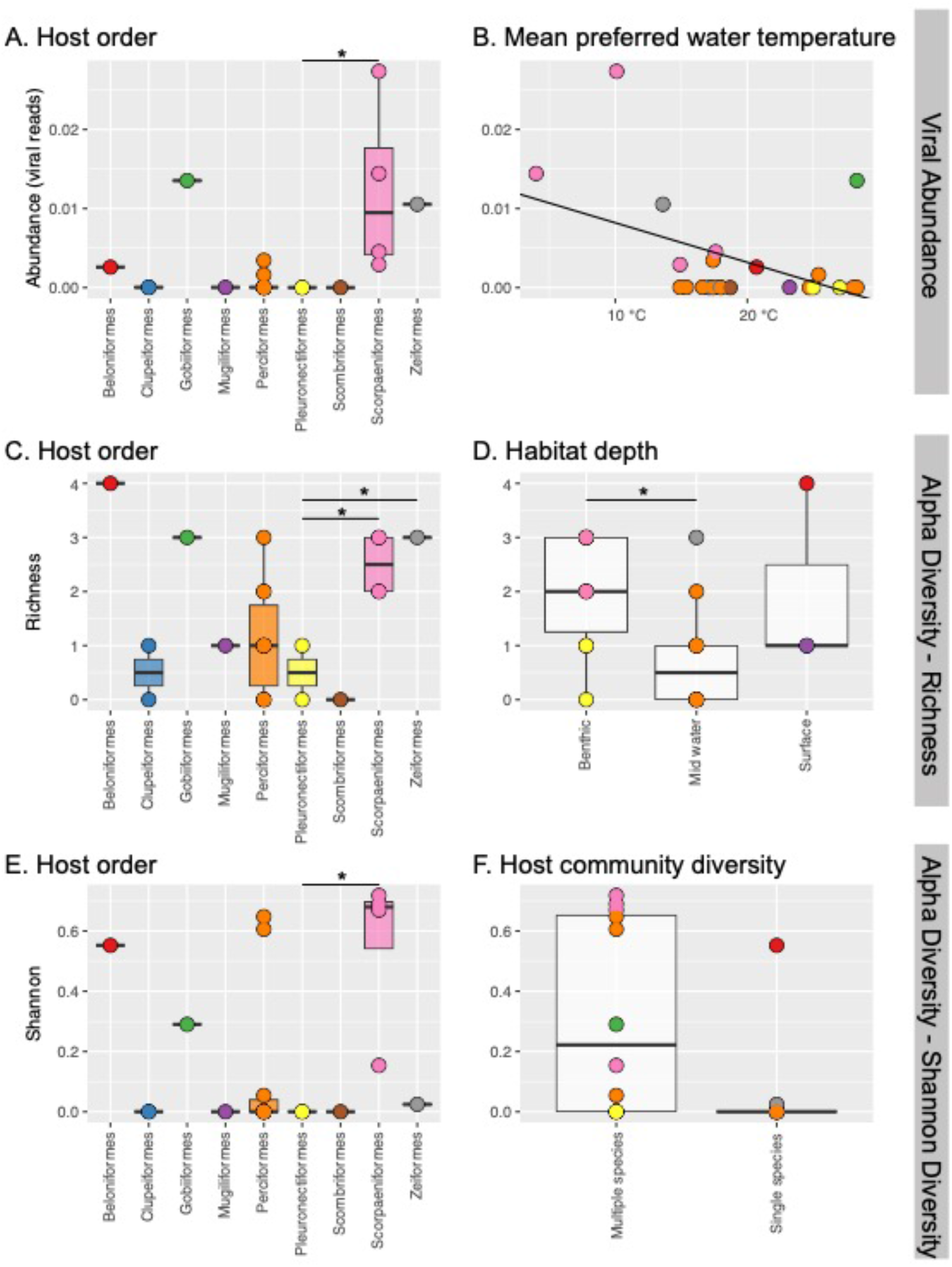
Significant explanatory variables in generalized linear models (GLM) for viral abundance and two measures of alpha diversity. Viral abundance is best explained by (A) fish host order and (B) mean preferred water temperature. Alpha diversity is best explained by (C) host order and (D) preferred habitat (Observed Richness) and by (E) host order and (F) host community diversity (Shannon Diversity). Stars indicate significant differences between groups determined by posthoc Tukey tests. Points represent different fish species and are coloured by host order.

We applied two measures of alpha diversity to our sample set: observed richness, a count of the number of viral families, and Shannon diversity, which also incorporates abundance. Observed richness was best explained by fish order (χ^2^=22.839, df=8, p=3.8^-6^) and habitat depth (χ^2^=3.914, df=2, p=0.032), while Shannon diversity was best explained by fish order (χ^2^=0.96, df=8, p=0.016) and community diversity (χ^2^=0.41, df=1, p=0.05), with a larger Shannon diversity in multispecies communities compared with single species communities. As with viral abundance, there was a significant difference in alpha diversity between Scopaeniformes compared to Pleuronectiformes (Tukey Richness z=3.039, p=0.0495; Tukey Shannon z=2.845, p=0.05). Notably, in these data mid-water fish had decreased viral richness compared to benthic fish (Tukey z=-2.452, p=0.0338), and fish that reside in multispecies communities had a larger Shannon diversity compared to single species communities (χ^2^=0.17089, df=1, p=0.05) (Figure 3). Our analysis also revealed that fish order (R^2^=0.57215, p=0.003), swimming behaviour (R^2^=0.09904, p=0.005), climate (R^2^=0.13315, p=0.012) and mean preferred water temperature (R^2^=0.1005, p=0.05) were significant predictors of beta diversity.

Importantly, we repeated the above analysis on the factors associated with virome composition on those viruses (n = 22) that likely infected hosts other than fish. Because we can assume that these viruses do not replicate in fish (for example, because they are related to host diet), and hence should be not shaped by aspects of fish biology and ecology, this analysis effectively constitutes an internal negative control. Indeed, this analysis revealed no association between virome composition and host ecological traits (viral abundance: p=0.0; alpha diversity: p=0.3; Shannon diversity: p=0.9; and beta diversity: p=0.3), thereby adding weight to the biological associations described above in the fish viruses.

## Discussion

The metagenomic revolution is enabling us to uncover more of a largely unknown virosphere. Here, we utilised mNGS to identify new viruses associated with fish, characterising the viromes of 23 species of marine fish that spanned nine taxonomic orders and identifying 47 novel viruses spanning 22 different virus families. This included 25 new vertebrate-associated viruses and a further 22 viruses associated with protozoans, plants, arthropods and fungi. Interestingly, the novel viruses included the first fish virus in the *Matonaviridae* that are the closest phylogenetic relatives of the mammalian rubella viruses. We also used these data to provide an initial assessment of how aspects of host biology might impact virus diversity and evolution. Although our study was limited to 23 fish species, on these data we found that host phylogeny (taxonomy) was strongly associated with the composition of fish viromes. We also identified several other host traits that were also associated with virus abundance and/or diversity, particularly preferred mean water temperature, climate, habitat depth, community diversity and whether fish swim in schools or are solitary. That these traits were not correlated with the composition of diet and microbiome-associated viruses that do not actively replicate in fish suggests that the patterns observed in marine fish are real, although it will clearly be important to test these initial conclusions using larger numbers of fish species sampled from a diverse set of environments.

Many of the viruses identified in this study were phylogenetically related to other, recently discovered, viruses of fish (Dill et al 2016, Geoghegan et al 2018a, Lauber et al 2017, Shi et al 2018a). However, there were some notable exceptions. Tiger flathead matonavirus represents the only fish viral species in the *Matonaviridae* and forms a distinct clade with a rubivirus discovered in a Chinese water snake. The discovery of this phylogenetically distinct fish virus tentatively suggests the possibility of a fish host origin for this family, although it is clear that confirmation will require the sampling of a far wider set of hosts. Indeed, it is notable that additional rubella-like viruses have recently been identified in a range of mammalian hosts, including bats (Bennett et al. 2020). A fish origin might also be the case for other virus families such as the *Hantaviridae* and *Filoviridae*, as the fish viruses in these families often fall basal to viruses in other vertebrate hosts such as birds and mammals (also see Shi et al 2018a). In contrast, in some other virus families such as the *Astroviridae, Picornaviridae, Flaviviridae* and *Rhabdoviridae*, viruses associated with fish are distributed throughout the phylogeny suggestive of a past history of common host-jumping. Regardless, available data suggests that fish viruses harbour more phylogenetic diversity than the better studied mammalian and avian viruses within these families. It is also clear that the discovery of novel viruses in fish has expanded our knowledge of the diversity, evolutionary history and host range of RNA viruses in general.

Although there is often a clear phylogenetic division between those viruses likely to infect fish and those associated with diet or microbiome, in some cases this separation can be nuanced. For instance, although totiviruses were thought to only infect unicellular fungi, their known host range has now expanded to include arthropods and fish (Mikalsen et al. 2016, Mor et al. 2016, Lovoll et al. 2010). In particular, piscine myocarditis virus is a totivirus shown by *in situ* hybridisation to infect Atlantic salmon and is associated with cardiomyopathy syndrome in salmon (Haugland et al. 2011). Similarly, viruses within the *Narnaviridae* are widespread in fungi, and have now been extended to include both invertebrates (Shi et al. 2016) and protist (Charon et al. 2019). Due to their phylogenetic position, we assume the narna-like viruses identified here are associated with fungal parasites in these samples.

As well as identifying new viruses, we sought to provisionally identify associations between host traits and the overall composition of fish viruses, although this analysis was clearly limited by the available sample size. A notable observation was that fish virome composition, reflected in measures of viral richness, abundance and diversity, is most impacted by the phylogenetic relationships (i.e. taxonomy) of the host in question. This in turn suggests that fish viruses might have co-diverged with fish hosts over evolutionary time-scales, a pattern supported by the general relationship between vertebrate host class and virus phylogeny observed for RNA viruses as a whole (Shi et al 2018a). However, it is also clear that cross-species is also a common occurrence in virus evolution (Geoghegan et al 2017). Indeed, it is possible that the strong association of host taxonomy and virome composition in some cases reflects preferential host switching among fish species (otherwise known as the ‘phylogenetic distance effect’; Longdon et al 2014), perhaps because viruses spread more often between phylogenetically closely related hosts due to the use of similar cell receptors (Charleston and Robertson 2002). These competing theories could be tested by more detailed co-phylogenetic comparisons among fish species that exhibit no ecological overlap thereby precluding cross-species transmission.

Our analysis also provided some evidence that virus abundance was negatively associated with the preferred water temperature of the fish species in question. Specifically, viruses were more abundant in fish that preferred cooler temperatures compared to those that prefer warmer temperatures. In this context it is noteworthy that virus transmission and disease outbreaks have been shown to be influenced by temperature and seasonality in farmed fish (Crane and Hyatt 2011). Moreover, for some viruses, host mortality is water temperature-dependent. For example, a highly infectious disease in fish, nervous necrosis virus, is more pathogenic at higher temperatures (Toffan et al 2016), while infectious hematopoietic necrosis virus, which causes disease in salmonid fish such as trout and salmon, causes mortality only at low temperatures (Dixon et al 2016). As the oceans continue to warm, it is crucial to understand the impact of increased temperatures on both marine life and virus evolution and emergence, especially as it is projected that outbreaks of marine diseases are likely to increase in frequency and severity (Dallas and Drake 2016, Karvonen et al 2010).

Also of note was that on these data, fish living in diverse, multi-fish species communities harboured more diverse viromes at a higher abundance than fish that live in less diverse, single-species communities. Previously, host community diversity has been hypothesised to lead to a decrease in infectious disease risk through the theory of the ‘dilution effect’ (Schmidt and Ostfeld, 2001). This theory views an increase in host species’ community diversity as likely to reduce disease risk, because encounter rates among preferred hosts are decreased, and both experimental and field studies have shown this phenomenon to occur across many host systems, particularly those involving vector-borne disease (Keesing et al 2006, LoGiudice et al 2003, Ostfeld and Keesing 2012). Although it might be reasonable to assume that increased virus abundance and diversity is directly correlated with disease risk, the association between host community diversity with that of virus diversity and abundance has not previously been tested. Our results, although preliminary, indicated that high multi-species community diversity in fish may be associated with increased virus diversity and abundance. It is possible that elevated community diversity in fish simply increases the total number of hosts in the system, in turn increasing viral diversity, particularly since host jumping appears to be common in fish viruses (Geoghegan et al 2018a).

Finally, it is noteworthy that since these fish species were market-bought rather than being directly sampled during fishing trips (with the exception of the pygmy goby), it is possible that viruses with short durations of infection were not detected. In addition, the relatively small number of individuals sampled here, and that samples were necessarily pooled to aid virus discovery, unavoidably limits some of the conclusions drawn. In particular, the host traits summarised here, such as life span, were taken at the overall species level rather than for the individuals sampled. It is therefore important to broaden sampling of fish and their viruses both geographically and seasonally, and include phenotypic data for the individuals sampled. This notwithstanding, our data again shows that fish harbour a very large number of diverse viruses (Shi et al. 2018; Lauber et al, 2017). Indeed, even the pygmy goby, one of the shortest-lived vertebrates on earth that lives for a maximum of 59 days on the reef (Depczynski and Bellwood 2005), harboured novel viruses that were assigned to three distinct virus families.

The new viruses discovered here greatly expand our knowledge of the evolutionary history of many virus families, particularly those with RNA genomes, with viruses identified in fish species that span highly diverse taxonomic orders. More broadly, the use of metagenomics coupled with a diverse multi-host, tractable system such as fish has the potential to reveal how host factors can shape the composition of viromes and that might ultimately lead to cross-species transmission and virus emergence.

## Data Availability

All sequence reads generated in this project are available under the NCBI Short Read Archive (SRA) under BioProject PRJNA637122 and all consensus virus genetic sequences have been deposited in GenBank under accession MT579871-MT579895

## Acknowledgements

We thank the New South Wales Department of Primary Industries for help sourcing fish samples. We thank efishalbum.com for fish images in Figure 1, which were used with permission. ECH and DRB are funded by ARC Australian Laureate Fellowships (FL170100022 and FL190100062, respectively). This work was partly funded by ARC Discovery grant DP200102351 to ECH and JLG and a Macquarie University Grant awarded to JLG.

## Supplementary Information

**SI Table 1**. Fish species sampled and the host features used in this analysis, with the latter obtained from fishbase.org. These features comprised fish taxonomic order, swimming behaviour (i.e. solitary or schooling fish), preferred climate, mean preferred water temperature, host community diversity (i.e. multi- or single-species community), average species length, maximum life span, trophic level and habitat depth.

**SI Table 2**. Amino acid identity, contig length and relative frequency of the viruses identified in this study. This does not include viruses described in Geoghegan et al (2018a).

**SI Table 3**. Accession numbers of viruses used to construct virus phylogenetic trees in Figure 2.

**SI Figure 1**. Graphical representation of virus families shared between fish species in this study. Fish species are shown in grey at the edges. The coloured connecting lines illustrate cases where viruses are present in both hosts for (A) viruses associated with fish hosts and (B) viruses likely associated with non-vertebrate host taxa (arthropods, fungi, plants, and protozoa). Caution must be met while interpretating this figure since virus abundance is not visually displayed. For example, sequence reads of reovirus were detected in the sand whiting at very low abundance (A) and this was insufficient to be included in the phylogenetic analysis shown in Figure 2. In addition, connecting lines depict hosts that share viruses from the same family and do not represent direct virus transmission between hosts. For both (A) and (B) only those fish species carrying viruses are shown.

**SI Figure 2**. Maximum likelihood phylogenetic trees depicting relationships among newly identified viruses associated with non-fish hosts that are members of the *Totiviridae, Narnaviridae* and *Partitiviridae*. Orange-coloured tips indicate the novel viruses within each family, while the relevant clades are highlighted with green-shaded boxes. Highly supported clades corresponding to SH-aLRT ≥ 80% and UFboot ≥ 95% are shown with white circles at the tree nodes. The number of amino acid substitutions per site is represented with a scale bar below each tree. The non-vertebrate host taxa (arthropods, fungi, plants, and protozoa) associated to each virus family are shown in the inset.

**SI Figure 3**. Maximum likelihood phylogenetic trees depicting relationships among newly identified viruses associated with non-fish hosts that are members of the families *Solemoviridae* and *Tombusviridae*. Orange-coloured tips indicate the novel viruses within each family, while the relevant clades are highlighted with purple-shaded boxes. Highly supported clades corresponding to SH-aLRT ≥ 80% and UFboot ≥ 95% are shown with white circles at the tree nodes. The number of amino acid substitutions per site is represented with a scale bar below each tree. The non-vertebrate host taxa (arthropods, fungi, plants, and protozoa) associated to each virus family are shown in the inset.

**SI Figure 4**. Maximum likelihood phylogenetic trees depicting relationships among newly identified viruses associated with non-fish hosts that are members of the families *Hepeviridae, Chuviridae, Nodaviridae, Iflaviridae, Dicistroviridae* and *Picornaviridae*. Orange-coloured tips indicate the novel viruses within each family, while the relevant clades are highlighted with blue-shaded boxes. Highly supported clades corresponding to SH-aLRT ≥ 80% and UFboot ≥ 95% are shown with white circles at the tree nodes. The number of amino acid substitutions per site is represented with a scale bar below each tree. The non-vertebrate host taxa (arthropods, fungi, plants, and protozoa) associated to each virus family are shown in the inset.

