## Supplementary figures and images for "Virome composition in marine fish revealed by meta-transcriptomics"

### Supplementary Figure 1

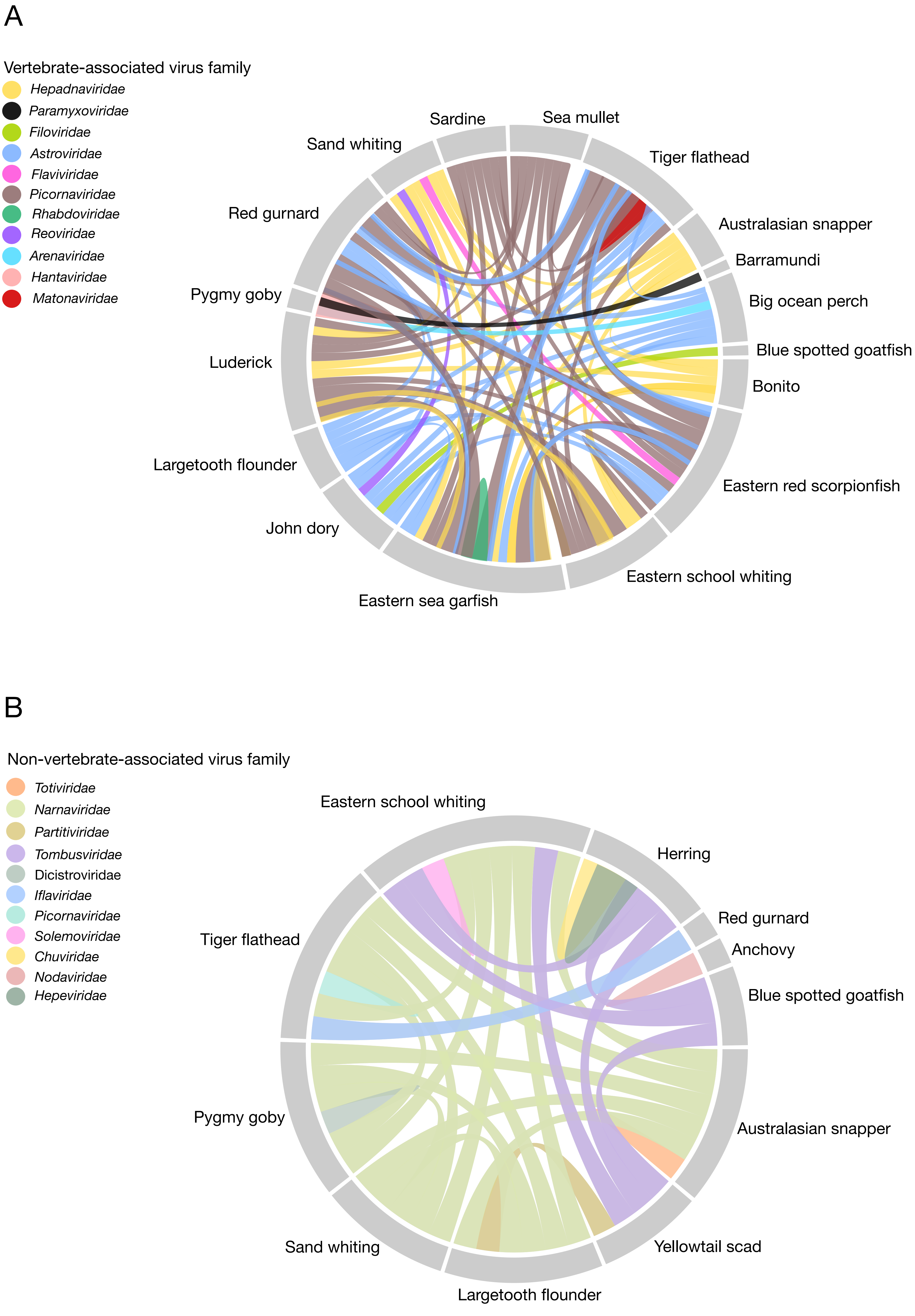

### Supplementary Figure 2

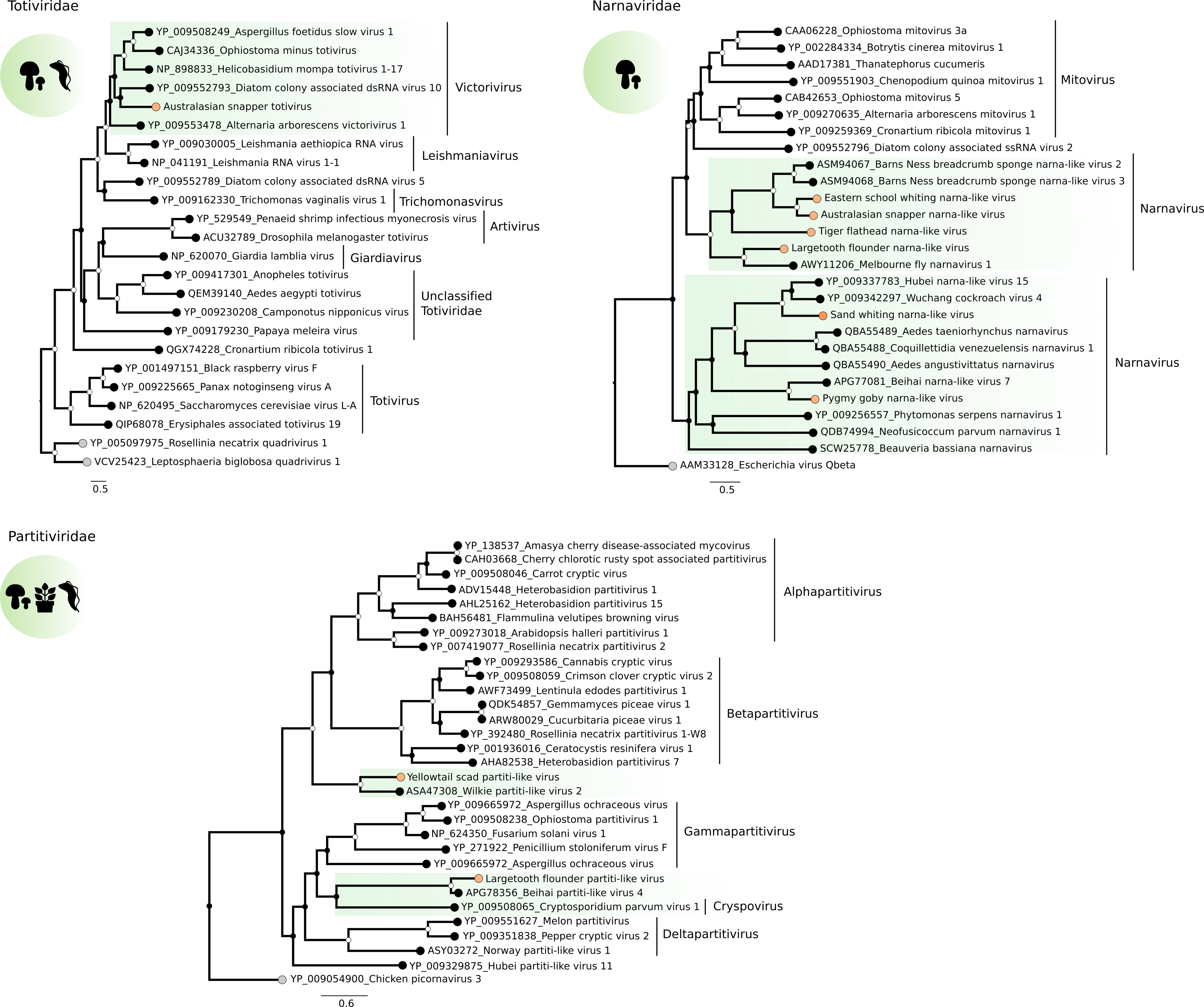

### Supplementary Figure 3

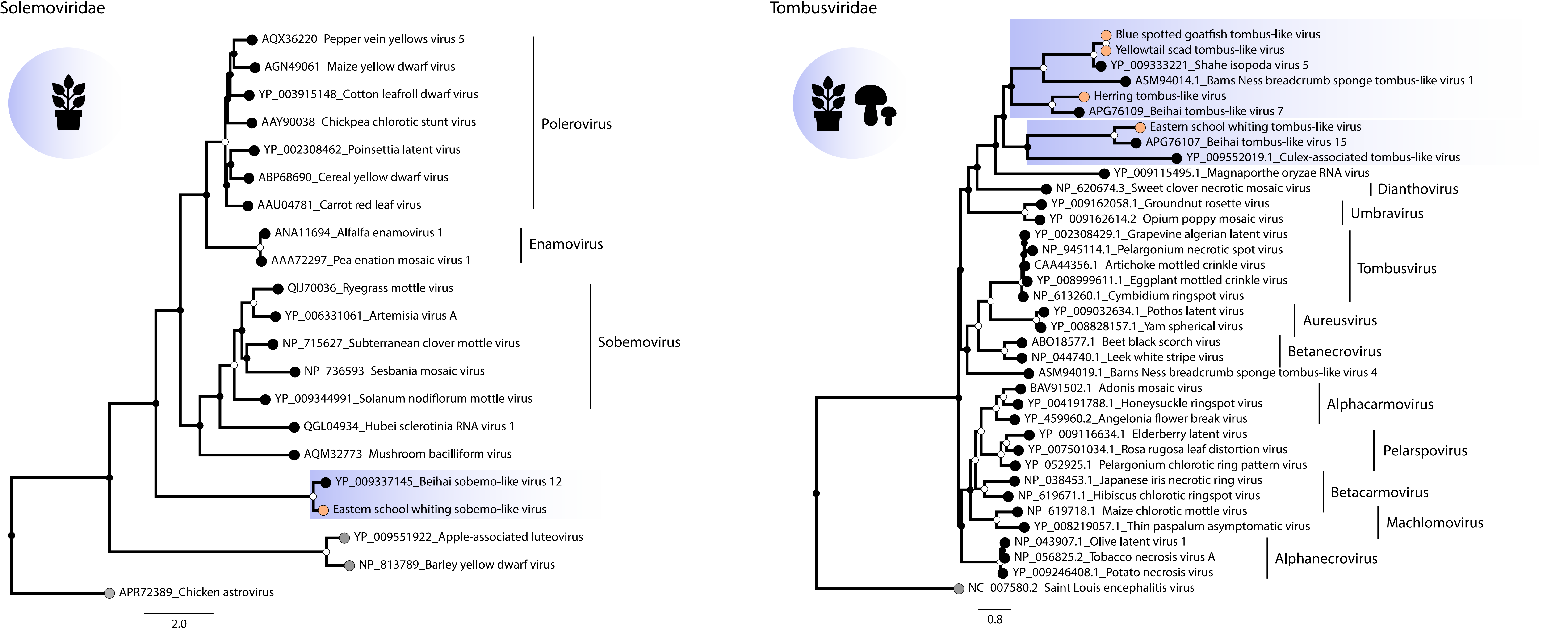

### Supplementary Figure 4

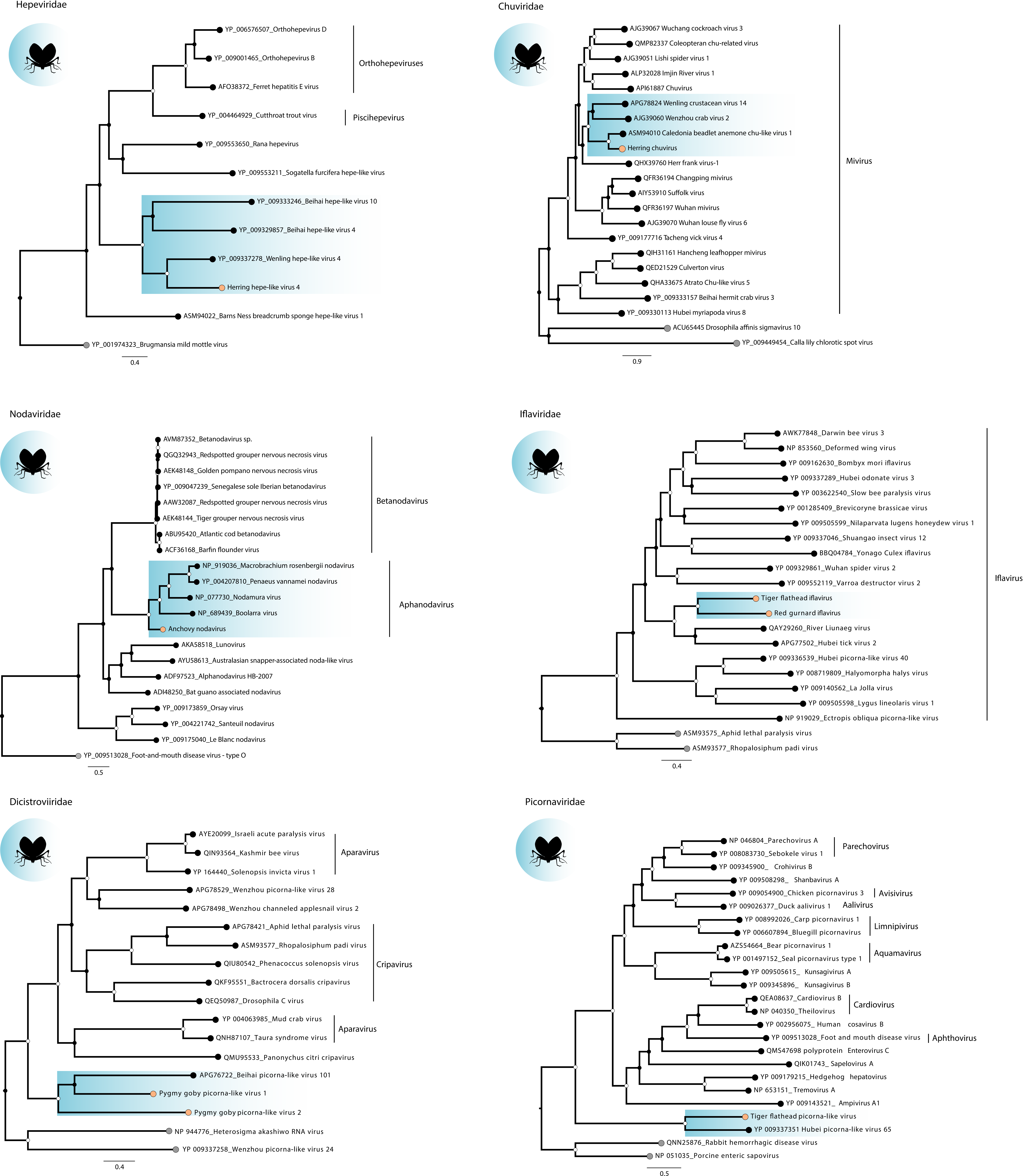
