## Supplementary Table 1 for "Virome composition in marine fish revealed by meta-transcriptomics"

| **Study** | **Fish** | **Species** | **Order** | **Swimming behaviour** | **Common**  **length (cm)** | **Trophic level** | **Depth range** | **Habitat** | **Community**  **diversity** | **Mean preferred**  **temperature (Celsius)** | **Climate** | **Max life span (years)** |
| --- | --- | --- | --- | --- | --- | --- | --- | --- | --- | --- | --- | --- |
| Geoghegan et al. 2018 | Australasian snapper | *Pagrus auratus* | Perciformes | both | 40 | 3.6 | 0 - 200 m | Mid | single/multi (age dependent) | 17.4 | Subtropical | 54 |
| Geoghegan et al. 2018 | Eastern red scorpionfish | *Scorpaena cardinalis* | Scorpaeniformes | Solitary | 47 | 3.5 | 0 - 154 m | Benthic | multi | 17.6 | Subtropical | 11 |
| Geoghegan et al. 2018 | Eastern sea garfish | *Hyporhamphus australis* | Beloniformes | Schooling | 29 | 2.6 | 0 - 20 m | Surface | single | 20.7 | Subtropical | 10 |
| Geoghegan et al. 2018 | Largetooth flounder | *Pseudorhombus arsius* | Pleuronectiformes | Solitary | 30 | 4.2 | 0 - 200 m | Benthic | multi | 27 | Tropical | 4 |
| This study | Bigeye ocean perch | *Helicolenus barathri* | Scorpaeniformes | Solitary | 40 | 4 | 285 - 739 m | Benthic |  | 4 | Temperate | 60 |
| This study | Red gurnard | *Chelidonichthys cuculus* | Scorpaeniformes | Solitary | 27.6 | 3.8 | 30 - 250 m | Benthic | multi | 10.1 | Temperate | 21 |
| This study | John Dory | *Zeus faber* | Zeiformes | Solitary | 40 | 4.5 | 5 - 400 m | Benthic - Mid | Single | 13.6 | Temperate | 12 |
| This study | Bonito | *Sarda australis* | Perciformes | Schooling | 45 | 4.5 | 0 - 30 m | Surface - Mid | Single | 14.9 | Temperate | 9 |
| This study | Tiger flathead | *Neoplatycephalus richardsoni* | Scorpaeniformes | Solitary | 45 | 3.9 | 10 - 400 m | Benthic | multi | 14.9 | Temperate | 15 |
| This study | Eastern school whiting | *Sillago flindersi* | Perciformes | Schooling | 25 | 3.3 | 0 - 60 m | Benthic | multi | 15.4 | Temperate | 7 |
| This study | Luderick | *Girella tricuspidata* | Perciformes | both | 35 | 2.1 | 0 - 20 m | Benthic - Mid | multi | 16.6 | Temperate | 11 |
| This study | Anchovy | *Engraulis australis* | Clupeiformes | Schooling | 12 | 3 | 31 - 70 m | Surface - Mid | single | 17.1 | Subtropical | 6 |
| This study | Herring | *Arripis georgianus* | Perciformes | Schooling | 20 | 4.3 | 0-50 m | Surface - Mid | single | 17.4 | Subtropical | 10 |
| This study | Sardine | *Sardinops sagax* | Clupeiformes | Schooling | 18 | 2.8 | 0 - 200 m | Surface | single | 17.9 | Subtropical | 9 |
| This study | Blue spotted goatfish | *Upeneichthys vlamingii* | Perciformes | Solitary | 35 | 3.5 | 5 - 100 m | Benthic | multi | 18 | Subtropical | 5 |
| This study | Blue mackerel | *Scomber australasicus* | Scombriformes | Schooling | 30 | 4.2 | 87-200 m | Surface - Mid | single | 18.7 | Subtropical | 8 |
| This study | Sea Mullet | *Mugil cephalus* | Mugiliformes | Schooling | 50 | 2.5 | 0 - 120 m | Surface | single | 23.2 | Subtropical | 15 |
| This study | Yellowfin bream | *Acanthopagrus australis* | Perciformes | Schooling | 40 | 3.1 | 0-50 m | Mid | single | 24.7 | Subtropical | 20 |
| This study | Smalltooth flounder | *Pseudorhombus jenynsii* | Pleuronectiformes | Solitary | 22.5 | 3.5 | 22 - 100 m | Benthic | multi | 25 | Tropical | 4 |
| This study | Sand whiting | *Sillago ciliata* | Perciformes | Solitary | 25 | 3.2 | 20 - 22 m | Benthic | multi | 25.4 | Tropical | 22 |
| This study | Yellowtail scad | *Trachurus novaezelandiae* | Perciformes | Schooling | 26 | 4.2 | 1 - 80 m | Surface - Mid | single | 28 | Tropical | 28 |
| This study | Barramundi | *Lates calcarifer* | Perciformes | both | 150 | 3.8 | 10 - 40 m | Mid | single/multi (age dependent) | 28.3 | Tropical | 35 |
| This study | Pygmy goby | *Eviota zebrina* | Gobiiformes | Solitary | 2 | 3.1 | 0 - 28 m | Benthic | multi | 28.3 | Tropical | 0.16 |
