## Supplementary Table 2 for "Virome composition in marine fish revealed by meta-transcriptomics"

| **Host** | **Virus** | **Virus family** | **Method of detection** | **Segment/gene and contig length (nt)** | **Standardised abundance** | **Closest Genbank match** | **Amino acid identity match (%)** |
| --- | --- | --- | --- | --- | --- | --- | --- |
| Pygmy goby | Pygmy goby hantavirus | Hantaviridae | Sequence similarity | Nucleoprotein: 534 | 2.34E-06 | Wenling yellow goosefish hantavirus (AVM87661.1) | 34.97 |
| Barramundi | Barramundi paramyxovirus | Paramyxoviridae | Sequence similarity | L protein (RdRp): 714 | 2.33638E-07 | Wenzhou pacific spadenose shark paramyxovirus (AVM87362.1) | 50.19 |
| Pygmy goby | Pygmy goby paramyxovirus | Paramyxoviridae | Sequence similarity | L protein (RdRp): 777 | 8.45716E-06 | Wenzhou pacific spadenose shark paramyxovirus (AVM87362.1) | 45.74 |
| Pygmy goby | Pygmy goby arenavirus | Arenaviridae | Sequence similarity | L segment (RdRp): 7089 | 0.000125538 | Wenling frogfish arenavirus 1 (YP_009551555.1) | 31.93 |
| Big eyed perch | Big eyed perch arenavirus | Arenaviridae | Sequence similarity | L segment (RdRp): 2919 | 5.13779E-06 | Wenling frogfish arenavirus 2 (AVM87649.1) | 58.25 |
| Sand whiting | Sand whiting flavivirus | Flaviviridae | Sequence similarity | Polyprotein (NS3): 258 | 1.61395E-05 | Guangxi houndshark hepacivirus (AVM87256.1) | 59.52 |
| John Dory | John Dory filovirus | Filoviridae | Sequence similarity | L protein (RdRp): 208 | 4.08946E-08 | Wenling thamnaconus septentrionalis filovirus (AVM87247.1) | 40.3 |
| Blue spotted goatfish | Blue spotted goatfish filovirus | Filoviridae | Sequence similarity | L protein (RdRp): 206 | 2.39808E-08 | Wenling thamnaconus septentrionalis filovirus (AVM87247.1) | 56.06 |
| Bonito | Bonito hapadnavirus | Hepadnaviridae | Sequence similarity | Polymerase: 893 | 3.34632E-07 | Rockfish nackednavirus (AZP02119.1) | 44.23 |
| Ludrick | Ludrick hapadnavirus | Hepadnaviridae | Sequence similarity | Polymerase: 493 | 1.612E-07 | Eastern sea garfish hepatitis B virus (AYU58612.1) | 31.73 |
| Eastern school whiting | Eastern school whiting hapadnavirus | Hepadnaviridae | Sequence similarity | Polymerase: 583 | 2.62634E-07 | African cichlid hepadnavirus (ANN02854.1) | 46.11 |
| Sand whiting | Sand whiting hapadnavirus | Hepadnaviridae | Sequence similarity | Polymerase: 933 | 1.61395E-05 | Bluegill hepatitis B virus (YP_009259541.1) | 48.18 |
| Eastern sea garfish | Eastern sea garfish rhabdovirus | Rhabdoviridae | Protein structural similarity | RdRp: 726 | 3.43744E-07 | Fujian dimarhabdovirus (AVM87298.1) | 44.9 |
| Tiger flathead | Tiger flathead matonavirus | Matonaviridae | Sequence similarity | NS polyprotein (RdRp): 4773 | 1.76297E-05 | Guangdong chinese water snake rubivirus (AVM87614.1) | 34.86 |
| Ludrick | Ludrick picornavirus | Picornaviridae | Sequence similarity | RdRp: 493 | 2.99372E-07 | Wenzhou picorna-like virus 43 (YP_009337305.1) | 81.71 |
| Eastern school whiting | Eastern school whiting picornavirus | Picornaviridae | Sequence similarity | RdRp: 230 | 1.09431E-07 | Fathead minnow picornavirus (AHL16617.1) | 47.92 |
| Seamullet | Seamullet picornavirus | Picornaviridae | Sequence similarity | RdRp: 457 | 2.63494E-07 | Ljungan virus (BAV53294.1) | 36.22 |
| Sardine | Sardine picornavirus | Picornaviridae | Sequence similarity | RdRp: 385 | 5.85112E-07 | Wenling rattails picornavirus (AVM87441.1) | 48.41 |
| Tiger flathead | Tiger flathead picornavirus | Picornaviridae | Sequence similarity | RdRp: 6633 | 1.11846E-05 | Wuhan sharpbelly picornavirus 1 (AVM87439.1) | 33.94 |
| Red gurnard | Red gurnard picornavirus | Picornaviridae | Sequence similarity | RdRp: 6870 | 0.000126102 | Wuhan sharpbelly picornavirus 1 (AVM87439.1) | 33.4 |
| John Dory | John Dory reovirus | Reoviridae | Sequence similarity | Segment 1 (RdRp): 3834 | 0.000105017 | Piscine orthoreovirus (QCT85126.1) | 63.1 |
| John Dory | John Dory astrovirus | Astroviridae | Sequence similarity | ORF1a (RdRp): 348 | 3.27157E-07 | Wenling pterygotrigla hemisticta astrovirus (AVM87158.1) | 50 |
| Red gurnard | Red gurnard astrovirus | Astroviridae | Sequence similarity | ORF1a (RdRp): 1544 | 0.000147438 | Wenling lepidotrigla astrovirus (AVM87156.1) | 85.94 |
| Tiger flathead | Tiger flathead astrovirus | Astroviridae | Sequence similarity | ORF1a (RdRp): 522 | 2.97464E-07 | Wenling perciformes astrovirus 2 (AVM87489.1) | 62.8 |
| Big eyed perch | Big eyed perch astrovirus | Astroviridae | Sequence similarity | ORF1a (RdRp): 3102 | 0.000139145 | Wenling righteye flounders astrovirus (AVM87607.1) | 40.57 |
